# Multiple intersecting pathways are involved in the phosphorylation of CPEB1 to activate translation during mouse oocyte meiosis

**DOI:** 10.1101/2024.01.17.575938

**Authors:** Chisato Kunitomi, Mayra Romero, Enrico Maria Daldello, Karen Schindler, Marco Conti

## Abstract

The RNA-binding protein cytoplasmic polyadenylation element binding 1 (CPEB1) plays a fundamental role in the regulation of mRNA translation in oocytes. However, the nature of protein kinase cascades modulating the activity of CPEB1 is still a matter of controversy. Using genetic and pharmacological tools and detailed time courses, here we have reevaluated the relationship between CPEB1 phosphorylation and the activation of translation during mouse oocyte maturation. We show that both the CDK1/MAPK and AURKA/PLK1 pathways converge on the phosphorylation of CPEB1 during prometaphase. Only inactivation of the CDK1/MAPK pathway disrupts translation, while inactivation of either pathway leads to CPEB1 stabilization. However, stabilization of CPEB1 induced by inactivation of the AURKA/PLK1 does not affect translation, indicating that destabilization/degradation can be dissociated from translational activation. The accumulation of the endogenous CCNB1 protein closely recapitulates the translation data. These findings support the overarching hypothesis that the activation of translation in prometaphase in mouse oocytes relies on a CDK1-dependent CPEB1 phosphorylation, and this translational activation precedes CPEB1 destabilization.

## Introduction

Completion of meiosis and the generation of a female haploid gamete competent for fertilization are developmental processes essential for propagation of the species. In a female, meiosis initiates during fetal development when chromosomes replicate, and homologous recombination is initiated (Seydoux and Braun, 2006). At the time when oocytes have undergone chromosome pairing and homologous recombination, they arrest meiotic progression and enter a prolonged period of cell cycle quiescence termed dictyate (Gosden and Lee, 2010). This period of quiescence ends at the time when fully grown oocytes are enclosed in antral follicles and are ready to be ovulated. The resumption of the meiotic cell cycle is triggered by the LH surge followed by the relief of somatic paracrine repressive signal that maintains meiotic arrest (Conti et al., 2012, Conti and Franciosi, 2018).

Several signaling pathways are activated and necessary for oocyte reentry into the cell cycle. For many species, including mammals, it is established that a rapid decline in cAMP concentrations in the oocyte, triggered by activation of a cGMP-inhibited cAMP phosphodiesterase (PDE3A), is the initial signal for reentry into the cell cycle (Conti et al., 2012, Vaccari et al., 2009, Jaffe and Egbert, 2017, Norris et al., 2009, Mehlmann, 2005). This decline in second messenger results in the inactivation of protein kinase A (PKA) and dephosphorylation of a CDC25 phosphatase, which then translocates to the nucleus (Pirino et al., 2009, Oh et al., 2010). This activation and the simultaneous inhibition of the nuclear WEE2 kinase(Oh et al., 2010) cooperate in the activation of the CDK1/cyclin complex, or M-phase promoting factor (MPF), the master regulator of the cell cycle(Kishimoto, 2018). The stabilization of the active MPF complex in the nucleus leads to phosphorylations of protein substrates that are essential to induce nuclear envelope breakdown (GVBD), and chromosome condensation(Ubersax et al., 2003). These processes, together with the assembly of a spindle, are all steps necessary for the subsequent prophase-to-metaphase I (MI) transition (Holt et al., 2013).

In *Xenopus* oocytes, an early translation of *Mos* mRNA is followed by an activation of the MAPK pathway thus signaling oocyte meiotic reentry (Gotoh et al., 1995, Ferrell, 1999). This translation may be triggered by a cyclin independent activation of CDK1 through interaction with a non-cyclin protein (Kim and Richter, 2007). Positive feedbacks between MAPK and CDK1 have been described and cooperate at different times during *Xenopus* oocyte meiotic progression (Abrieu et al., 2001). *Mos* and *Erk* are essential to maintain the metaphase II (MII) arrest through stabilization of the cytostatic factor (CSF) (Kishimoto, 2003). Although the implication of *Mos/Erk* in MII arrest is also well established in mouse (Colledge et al., 1994), it is less clear the role of this signaling cascade during early meiotic cell cycle reentry. Reentry into meiosis occurs in mice null for MOS (Colledge et al., 1994) and when the downstream Erk1/Erk2 kinases have been genetically inactivated (Sha et al., 2017). However, several defects in spindle organization and localization have been detected (Sha et al., 2017).

Independent of CDK1, an additional signaling pathway is functional early in maturation prior to GVBD. Aurora kinase A (AURKA) is activated around microtubule-organizing centers (MTOCs) (Blengini et al., 2021) either by a change in its conformation through TPX2 binding or more broadly in the cytoplasm via phosphorylation by the kinase BORA (Joukov and De Nicolo, 2018). AURKA activation then phosphorylates PLK1 (Baran et al., 2016, Solc et al., 2015), and these steps are necessary to assemble the spindle. Small molecular inhibitors and genetic manipulation of AURKA document that this pathway is essential for spindle function and meiotic progression (Blengini et al., 2022). Finally, several reports have detected activation of the PI3 kinase and the downstream AKT pathway during early oocyte maturation (Kalous et al., 2006). Activation of this pathway may contribute to regulation of translation by promoting the assembly of a complex required for translation initiation (Susor et al., 2015).

Although these cascades of phosphorylation and activation/inactivation of kinases and phosphatases play pivotal roles in meiotic reentry, regulation of translation of mRNAs and synthesis of components critical for meiotic progression are also indispensable for meiotic reentry in *Xenopus* and for meiotic progression in mouse oocytes. In addition to translational regulation of *Mos*, cyclin synthesis is also critical to sustain CDK1 activity (Mendez and Richter, 2001, Conti and Kunitomi, 2024). The initial GVBD step in mouse oocytes occurs even in the presence of a protein synthesis inhibitor, suggesting that protein synthesis is dispensable for the initial CDK activation. However, the steady state levels of cyclins are the result of active turnover in quiescent oocytes indicating that translation of mRNAs and protein synthesis do have role early on during oocyte maturation (Reis et al., 2006, Marangos et al., 2007).

Although other RNA binding proteins such as MUSASHI may play a role in very early activation of *Mos* mRNA (Charlesworth et al., 2006), it is established that the RNA binding protein CPEB1 plays a crucial function in the translational regulations throughout maturation of the oocyte in both *Xenopus* and mice. In *Xenopus* and mouse oocytes, CPEB1 phosphorylation is required to assemble a complex that promotes polyadenylation of an mRNA and its translational activation (Richter and Lasko, 2011). Numerous studies have been published on the phosphorylation of CPEB1 and the consequent translational regulation in *Xenopus*. It was initially demonstrated that Aurora A/Eg2 kinase phosphorylates CPEB1 early (Mendez et al., 2000b, Mendez et al., 2000a). However, additional studies have questioned a role for this kinase in CPEB1 phosphorylation (Keady et al., 2007, Radford et al., 2008, Frank-Vaillant et al., 2000). Later during MI, CPEB1 receives additional phosphorylation by CDK1/ERK and these phosphorylation signal destabilization of CPEB1 (Mendez et al., 2002). This destabilization is thought to cause an activation of translation of mRNAs like *CcnB1* but not *Mos*. In addition to CDK1, a PLK1-mediated phosphorylation may also trigger degradation of CPEB1 (Setoyama et al., 2007). The kinases involved in phosphorylation of CPEB1 in mammalian oocytes is equally unclear. Initial studies reported data indicating that AURKA phosphorylates CPEB1 also in mouse oocytes (Hodgman et al., 2001). However, follow up studies using pharmacological inhibition concluded that AURKA is not the kinase involved in CPEB1 phosphorylation and translational activation in pig and mouse oocytes (Komrskova et al., 2014, Han et al., 2017).

More recent genetic studies using a null *Aurka* allele have reproposed a role for this kinase in translational regulation in mouse oocytes (Aboelenain and Schindler, 2021). As in *Xenopus*, CDK1 dependent phosphorylation of CPEB1 in mouse oocytes is likely responsible for its degradation around MI (Tay et al., 2000). Others have proposed that MAP kinase-dependent phosphorylation are the most relevant pathways signaling CPEB1 degradation (Sha et al., 2017). It is also proposed that destabilization is important for activation of translation (Sha et al., 2017). Therefore, as reported in *Xenopus*, partial degradation of CPEB1 activates translation (Zhang et al., 2018). Here, we have reassessed the different signaling pathways converging on CPEB1 phosphorylation and translational activation in mouse oocytes.

One aspect of translational regulation to consider is the timing of the different molecular events activated by the somatic signal that triggers oocyte maturation. A widely used *in vitro* model to study oocyte maturation relies on the decrease in cAMP by removing it from the culture medium (dbcAMP) or by washing out a phosphodiesterase (PDE) inhibitor such as IBMX, milrinone or cilostamide. Within one hour after washout, AURKA activation is detected at multiple MTOC foci (Solc et al., 2015), and CDK1 is activated first in the cytoplasm and then in the nucleus. These events are rapidly followed by GVBD (Oh et al., 2010). The timing of MAPK activation is less defined in mouse oocytes, with some reports detecting downstream ERK phosphorylation after 4 h (Su et al., 2002) while others reporting earlier activation (Cao et al., 2020). Messenger RNA translational activation or repression is detected right after GVBD around 2-3 h (Luong et al., 2020). Therefore, in defining the signaling pathways controlling translational activation, one should consider whether the timing in which each molecular event develops is compatible with the downstream translational activation. Here, we have investigated how different signaling pathways activated at the time of GVBD and meiotic reentry contribute to activation of translation.

## RESULTS

### Translational activation of CcnB1 and Mos mRNAs requires CPEB1

Given the pivotal roles of Cyclin B1 (CCNB1) and MOS in regulating meiotic cell cycle progression, several reports have investigated the transcriptional and post-transcriptional regulatory mechanisms governing the expression of these proteins. Previous studies explored the involvement of the RNA-binding protein CPEB1 in the translational activation of *Cyclin* and *Mos* mRNAs in mouse oocytes, using *Cpeb1* mRNA siRNA knockdown or mutagenesis of the 3’ untranslated region (UTR) of target mRNAs (Tay et al., 2000, Yang et al., 2017, Han et al., 2017). Here, we employed *in vivo* genetic manipulations to assess how CPEB1 is involved in the translational activation of *CcnB1* and *Mos* mRNAs. *Cyclin* and *Mos* reporter mRNAs, in which the 3’ UTR control the accumulation of the fluorescent protein YFP, were co-injected with *mCherry* reporter into quiescent prophase I-arrested (GV) mouse oocytes isolated from wild-type (WT) and CPEB1 oocyte-specific knockout (Cpeb1^fl/fl^Zp3-cre) mice (Fig. 1A). We then monitored reporter accumulation using time-lapse microscopy during oocyte maturation (Supplementary Fig. 1A-D). When assessing the changes in *Ccnb1* and *Mos* translation, an increase in rates during oocyte maturation was detected in oocytes from WT controls (Fig. 1B,C); however, translation rates for both were significantly decreased in oocytes derived from *Cpeb1^fl/fl^*Zp3-cre mice (Fig.1B,C) (FDR<0.0001). Upon examining the timing of reporter accumulation in more detail, an increase in translation rate became significant 2-3 hours after release from the cilostamide (PDE inhibitor) block (30 minutes-1 h after GVBD) in oocytes from WT mice (arrow (↓) in Fig. 1B,C). This timing is similar or slightly delayed in comparison with ribosome loading onto endogenous mRNAs encoding the two proteins (Luong et al., 2020).

**Figure 1.**
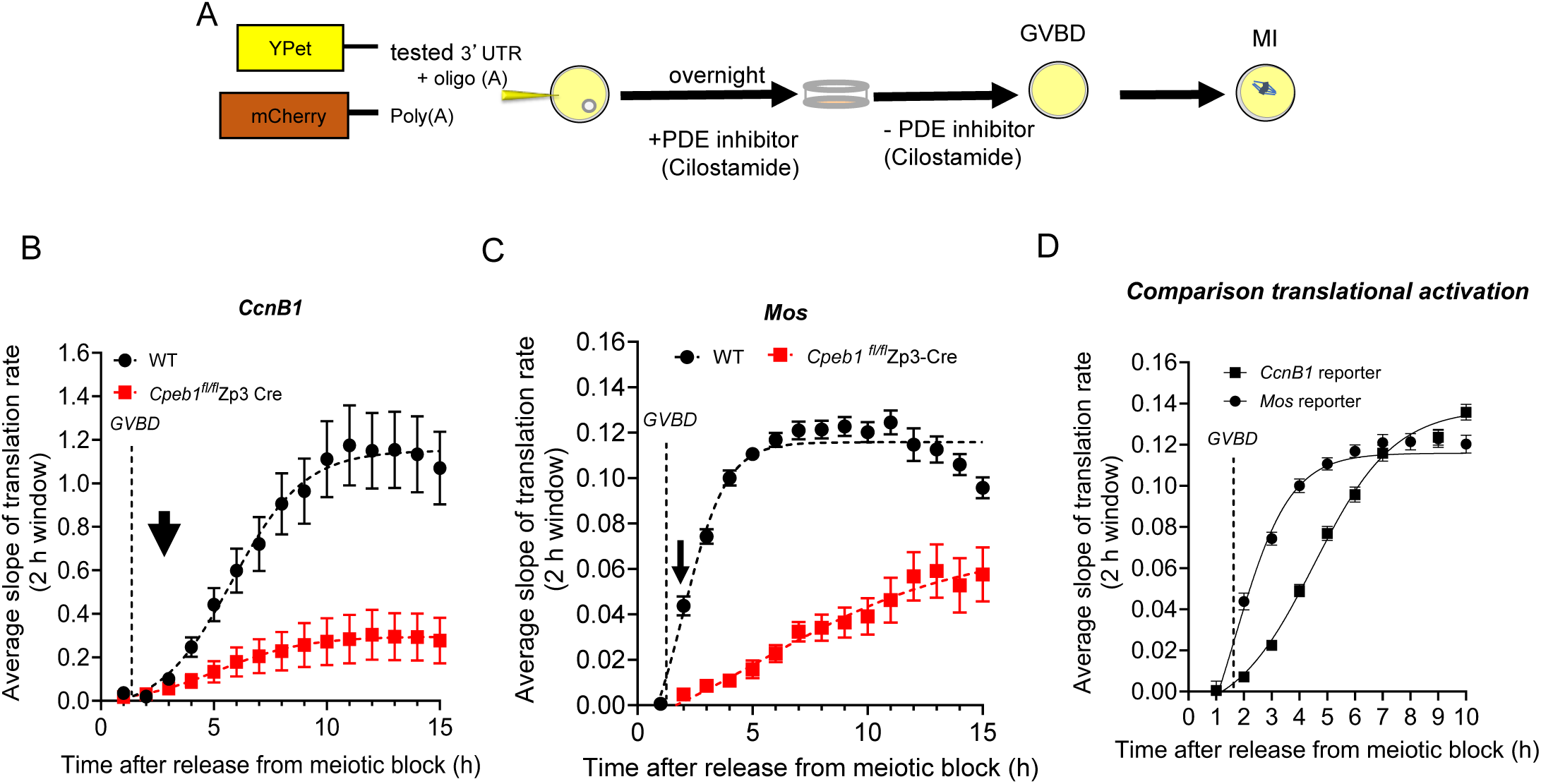
CPEB1 plays a major role in translational activation of *CcnB1* and *Mos* mRNAs. (A) Schematic representation of the YFP reporter assay. GV-arrested oocytes were injected with 5-10 pl of 12.5 ng/µl *CyclinB1 (Ccnb1)* or *Mos* reporter and 12.5 ng/µl polyadenylated mCherry. After overnight incubation with 1 µM cilostamide (PDE inhibitor), oocytes were released from the meiotic block. YFP and mCherry signals were recorded during maturation by time-lapse microscopy every 15 min for 20 h. The translation rate for each oocyte was calculated by linear regression of the reporter data using a 2-hour moving window and the mean ± SEM of all the oocytes from three independent experiments is plotted against incubation time. (B) Translation rates of *Ccnb1* reporter in wild type (WT) and *Cpeb1*^fl/fl^Zp3-cre oocytes during maturation (difference between the two groups: P<0.001 by nonparametric Mann-Whitney test). (C) Translation rate of *Mos* reporter in WT and *Cpeb1*^fl/fl^Zp3-cre oocytes during maturation (difference between the two groups: P<0.00001 by nonparametric Mann-Whitney test) (D) Comparison of the timing of *Ccnb1* and *Mos* translation rates in WT oocytes during maturation. The data are plotted as the mean ± SEM. Nuclear envelope breakdown (GVBD) timing (average 1.75 h) is shown in the graph. The arrow indicates the first point in the time course when a change in the rate of translation is significant. Significantly different at P<0.00001 using non parametric Mann-Whitney test.

Ribosome loading increased immediately after GVBD at the 2 h time point (Han et al., 2017, Luong et al., 2020). Of note, the timing of translational activation of these mRNAs after GVBD differs from that observed in *Xenopus* oocytes, where *Mos* mRNA translation is activated prior to GVBD, and *CcnB1* mRNA translation is delayed until well into prometaphase (Mendez et al., 2002, Pique et al., 2008). Despite activation of both occurring post-GVBD in mice, a difference in the timing of translation activation of the two reporters was detected. We found that translational activation of *Mos* mRNA preceded that of *CcnB1* (Fig. 1D), a difference reminiscent of what has been described in *Xenopus* oocytes.

When using oocytes from *Cpeb1^fl/fl^*Zp3-cre mice which we injected with a *Ccnb1* 3’ UTR reporter, we observed residual translational activation (Luong et al., 2020). This finding was confirmed and extended here where in different experiments over a period of 3 years we detected residual (21±6%) translational activation in oocytes with CPEB1-loss of function (Supplementary Fig. 1A-D). In agreement with our previous report (Luong et al., 2020), we detected 24±3% of CPEB1 protein remaining by western blot of extract of oocytes from a homozygous null background (Supplementary Fig. 1E), providing a possible explanation for the incomplete block of translational activation. Regardless of this incomplete removal of the protein, our data support the conclusion that CPEB1 plays a major role in the translation of both *CcnB1* and *Mos* mRNAs.

### Multiple pathways control phosphorylation of CPEB1 in oocytes

To delve further into the different molecular events necessary for translational activation during oocyte meiotic reentry, we reconsidered the different phosphorylation cascades reported to converge on CPEB1 and translational regulation.

To probe the time dimension of these events, we used both loss-of-function and small molecule pharmacological inhibition approaches to investigate the contribution of pathways leading to CPEB1 phosphorylation and translational activation. Although in previous reports we used a pharmacological strategy to manipulate the relevant kinases (Han et al., 2017, Luong et al., 2020), here we introduced two major modifications in the experimental protocol. First, we used culture conditions where oil was omitted during oocyte incubation, because the hydrophobicity of small molecule inhibitors used may partition to the oil phase (Gaudeline Rémillard Labrosse, 2023). Second, because inhibition of any of these pathways prevents oocyte progression to metaphase II (MII), all measurements were performed at 7-8 h of incubation, when the oocytes are still in metaphase I (MI). This latter change in experimental protocol should remove the confounding variable of oocytes being at different stages of the meiotic cell cycle depending on the treatment. Using these experimental settings, we focused on two major interconnected pathways converging on CPEB1: the AURKA/PLK1 and CDK1/MOS-ERK pathways.

To determine whether these pathways directly or indirectly converge onto the phosphorylation status of CPEB1, we assessed the shift in CPEB1 electrophoretic mobility on SDS-PAGE. A phosphoproteome analysis (Cheng et al., 2022) detected multiple CPEB1 phosphorylation sites in mouse oocytes (Supplementary Fig. 2A). These phosphosites fall within CDK1, MAPK, AURKA and PLK1 consensus sites. We and others have shown that multiple, stepwise changes in mobility and generation of different CPEB1 immunoreactive forms are detected during mouse oocyte maturation (Tay et al., 2000, Han et al., 2017, Yang et al., 2017). A minor shift in CPEB1 immunoreactivity during the first hour of oocyte culture was observed only under some electrophoretic conditions (Han et al., 2017), but a more significant mobility shift was consistently observed at 2 h. We refer to this CPEB1 species as the phosphorylated form (Supplementary Fig. 2B). This shift is followed by additional decreases in mobility between 3 and 5 h (Supplementary Fig. 2B). As in other reports (Mendez et al., 2002), we referred to this latter slowest migrating form at 4 h as the hyper-phosphorylated form.

When using this mobility readout to assess the effect of inhibition of the different signaling pathways, we found that the mobility of CPEB1 immunoreactive band is clearly affected in oocytes derived from oocyte-specific AURKA KO (conditional; cKO) mice (Fig. 2A). Specifically, the accumulation of the hyper-phosphorylated form is delayed and the ratio of phospho/hyper-phosphorylated forms at 4 h of meiotic maturation is inverted: WT oocytes accumulated more of the hyper-phosphorylated form whereas more of the phosphorylated form was present in AURKA cKO oocytes (Fig. 2B,C). The involvement of AURKA in generating this phosphorylation pattern was confirmed by inhibition of AURKA activity with a concentration of MLN8237 that preferentially block this kinase (Blengini et al., 2022) (Fig. 2A-C). This small molecule inhibitor mimicked the genetic loss-of-function, producing less of the hyper-phosphorylated form at 4 h (Fig. 2A-C). Pharmacological inhibition of PLK1 by using Bi2536, a kinase phosphorylated and activated by AURKA (Willems et al., 2018), produced even a more profound effect and the hyper-phosphorylated form was not detected at 4 h in some experiments (Fig. 2A-C). Control experiments where AURKA phosphorylation was monitored by western blotting confirmed a complete inhibition by these treatments (Supplementary Fig. 3B). Thus, and unlike our previous conclusions (Komrskova et al., 2014, Han et al., 2017), AURKA either directly or indirectly affects the phosphorylation state of CPEB1.

**Figure 2.**
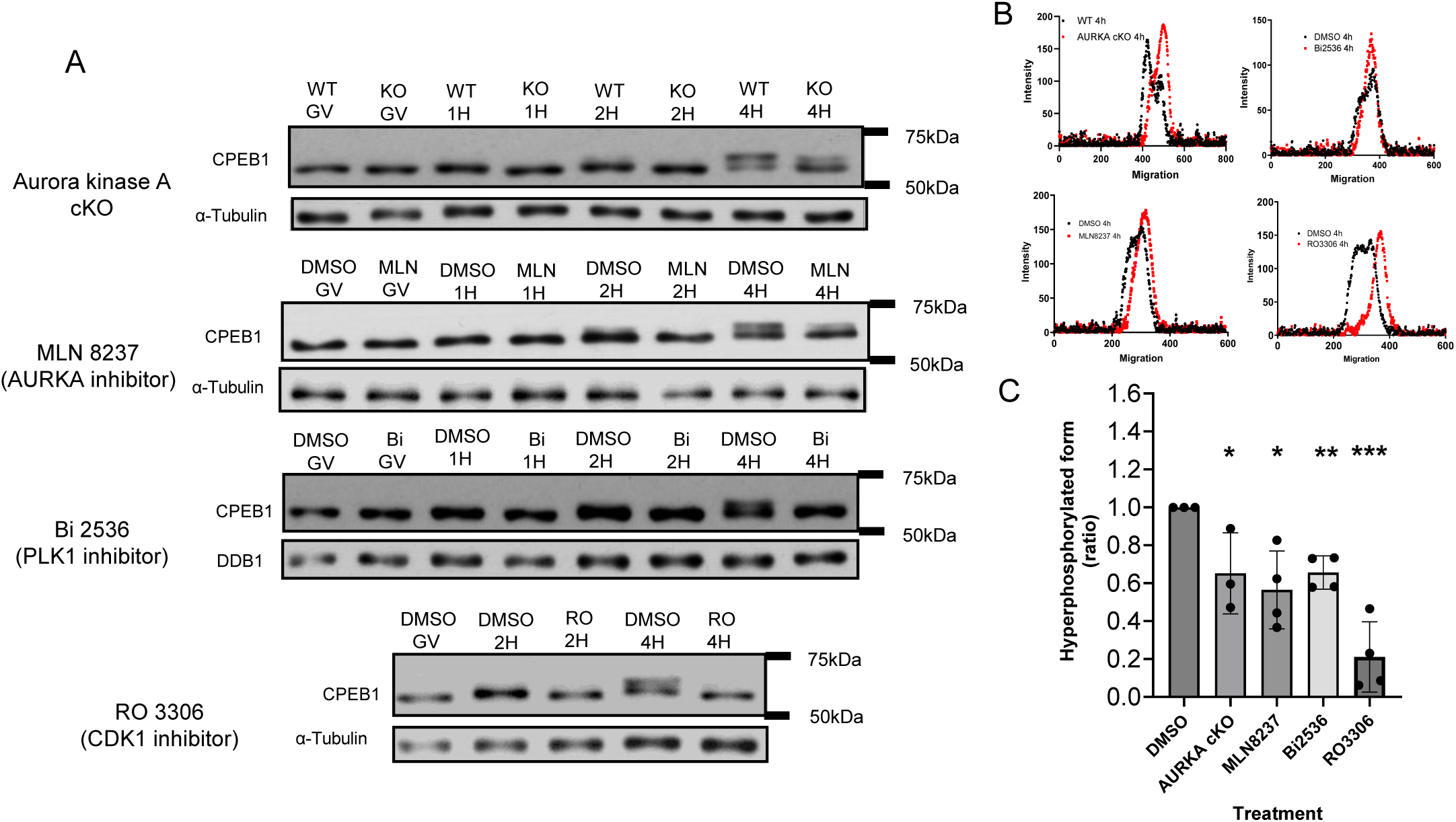
CPEB1 phosphorylation by multiple pathways. (A) Representative western blot images of the time course of CPEB1 phosphorylation with different treatments. Western blot analyses were conducted on lysates of 30 oocytes at the reported time points. Alpha F-Tubulin or DDB1 was used as a loading control. (B) The profile of band intensity at 4 h in control (black dots) and experimental (red squares) groups are plotted against the migration on the gel. (C) The integration of hyper-phosphorylated peak intensities was calculated and compared to DMSO. Each point is the ratio of experimental/DMSO control and represents a different experiment and a different biological replicate of the three or more experiments performed. Two-tailed unpaired Student’s test was used to evaluate the statistical significance between DMSO and each treatment (*P<0.05, **P<0.01, ***P<0.001). Note that, with the exception of the RO3306 profile, the abundance of the hyper-phosphorylated band in each treatment is over-estimated because not completely resolved from the phosphorylated band. WT: wild type, KO: knock out, GV: prophase I-arrested oocytes, AURKA: Aurora Kinase A.

Several CPEB1 phospho-peptides containing CDK1 consensus sites were identified by mass spec in a previous study (Supplementary Fig. 2A). Inhibition of CDK1 activity with the specific RO3306 inhibitor prevented the generation of both the phosphorylated and hyper-phosphorylated forms (Fig. 2A-C). To confirm the efficacy of treatments, we performed western blotting to monitor ERK phosphorylation and found that inhibition of CDK1 prevented ERK activation (Supplementary Fig. 3C).

Because AURKA/PLK1 and CDK1/MAPK pathways intersect at several steps through positive and negative feedbacks, we tested whether inhibition of AURKA signaling affects the activation of MAPK by assessing its phosphorylation state. When assessing GVBD timing by time lapse microscopy, a 30 min delay in oocyte GVBD was observed in AURKA cKO oocytes as well as after pharmacological inhibition of this pathway (Supplementary Fig. 3A), suggesting that altered phosphorylation around MTOCs delays CDK1 activation. However, in all cases no decreased MAPK phosphorylation could be detected by western blot of AURKA cKO oocytes, and in the MLN8237 and Bi2536 treatments which inhibit AURKA and PLK1, respectively (Fig. 4C, Supplementary Fig. 3C). Thus, it is unlikely that the altered CPEB1 phosphorylation that follows manipulation of AURKA signaling is due to altered CDK1 function. Note also that CDK1 inhibition completely blocked MAPK phosphorylation, a finding consistent with the view that CDK1 activation functions upstream of MAPK activation, likely through control of *Mos* mRNA translation (Supplementary Fig. 3C) or MOS protein stabilization (Castro et al., 2001, Martina Santoni, 2023).

In summary, these experiments indicate that timely phosphorylation of CPEB1 during prometaphase is dependent on both functional AURKA and CDK1 pathways.

### Inhibition of phosphorylation of CPEB1 causes stabilization of the protein and/or delayed degradation

Previous reports indicated that hyper-phosphorylation of CPEB1 is associated with destabilization of the protein (Mendez et al., 2002). It has been proposed that this posttranslational modification is a mechanism of translation activation because CPEB1 represses 3’UTR that contain multiple CPEs (Mendez et al., 2002, Sha et al., 2017). We then determined whether the manipulation of the above pathways also affect the stability of CPEB1 in MI (Fig. 3A,B). Both genetic and pharmacological inhibition of the AURKA/PLK1 pathway caused partial or complete stabilization of CPEB1 protein at 7 h (Fig. 3A,B). Although hyper-phosphorylation was still present in AURKA cKO oocytes and after MLN8237 treatment of WT oocytes at 7 h, this species was decreased in PLK1 inhibited oocytes (Fig. 3A). As mentioned, the loss of function of AURKA caused a 30-40 min delay in GVBD, and similar effects were detected with MLN8237 and Bi2536 treatments (Supplementary Fig. 3A). This short delay in reentry into the cell cycle did not impact the time course of degradation since the stabilization of CPEB1 after inhibition of PLK1 was detected at 7 h and even more at 8 h, a time when oocytes reached MI (Supplementary Fig. 4). Loss of both phosphorylated and hyper-phosphorylated forms, and CPEB1 stabilization in the unphosphorylated state is evident with inhibition of CDK1 (Fig. 3A). Stabilization was also observed with the MAPK inhibitors; however, the migration of the stabilized form corresponded to that of a hyper-phosphorylated CPEB1 (Fig. 3A,B). Taken together, these findings are consistent with the hypothesis that activation of both pathways is involved in CPEB1 hyper-phosphorylation and destabilization.

**Figure 3.**
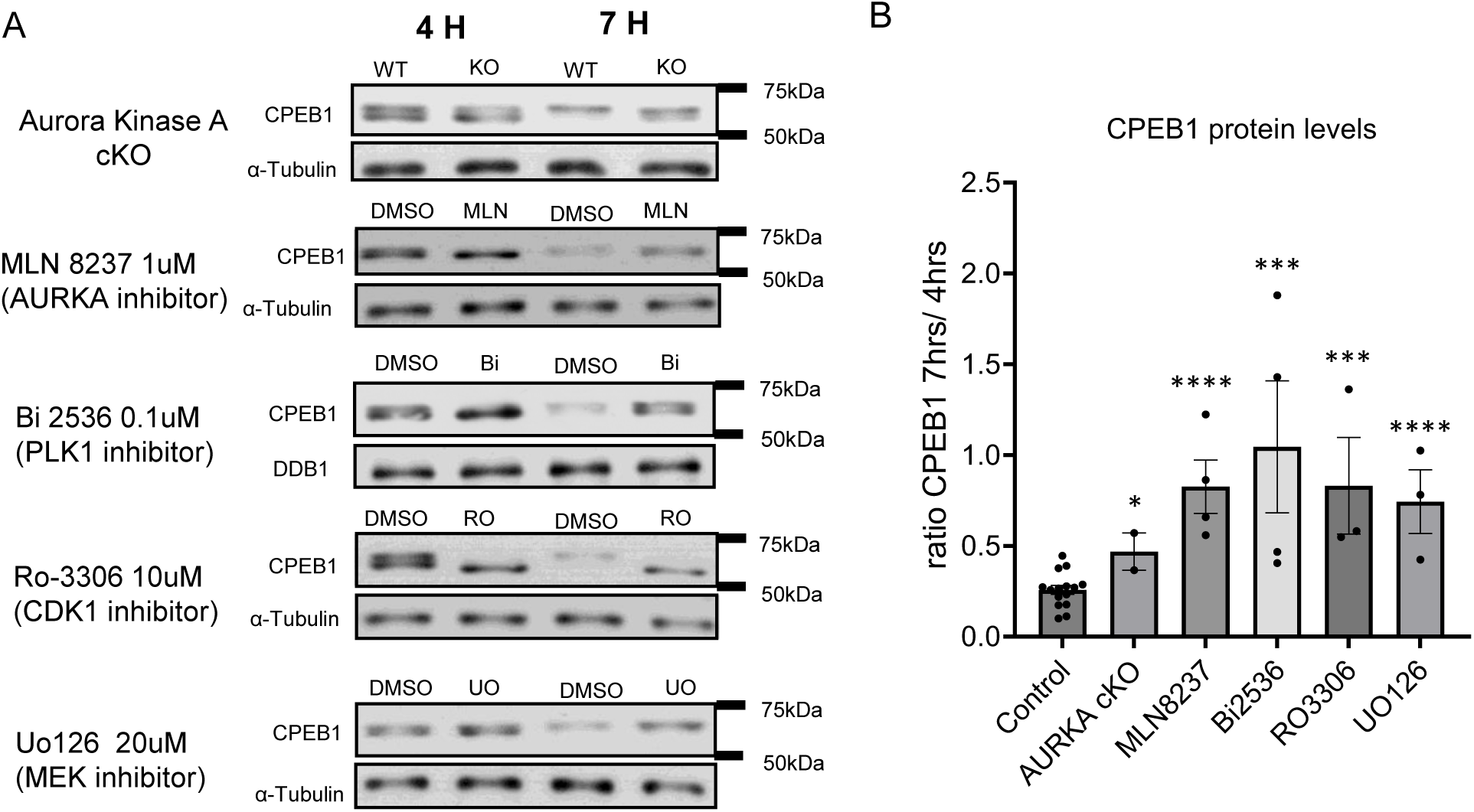
CPEB1 stabilization induced by the inhibition of different protein kinases. (A) Representative western blot images of the CPEB1 phosphorylation at 4 and 7 h with different treatments. Western blot analysis was conducted on lysates of 30 oocytes. Alpha-Tubulin or DDB1 was used as a loading control. (B) CPEB1 stabilization was calculated as the ratio of the CPEB1 immunoreactive signal between 7 h and 4 h. Each bar is the mean ± SEM of three or more independent biological replicates, except for the AURKA cKO group where the range of two experiments is plotted. Two-tailed unpaired Student’s test was used to evaluate the statistical significance between DMSO and each treatment (*P<0.05, ***P<0.001, ****P<0.0001). WT: wild type, KO: knock out, AURKA: Aurora Kinase A.

### Inhibition of the AURKA pathway does not compromise Ccnb1 mRNA translation and CCNB1 protein accumulation

Given the prominent function of CPEB1 in translational activation during oocyte maturation, we next investigated how interference with any of the above phosphorylations would affect translation of *CcnB1* mRNA, a prototypic target of this RNA-binding protein (RBP). The genetic ablation of AURKA had no significant effect on the translational activation of *Cyclin B1* reporter translation (Fig. 4A) and the rate of translation in the absence of AURKA activity at 6-8 h was not significantly different from WT controls (Fig. 4B). The absence of any effect when AURKA is absent was confirmed even when protracting the incubation for 15 h (Fig. 4A). Similarly, activation of translation in the presence of the AURKA or PLK1 (MLN8237 or Bi2536) inhibitors was not significantly reduced (Fig. 5A,B), even though a trend toward a decrease was detected with PLK1 inhibition. Conversely, CDK1 inhibition caused a profound decrease in translational activation when applied immediately after GVBD (Fig. 5C). The delayed treatment was necessary to allow oocytes to undergo GVBD. An 80% decrease in translation rate was observed at 8 h (Fig. 5C). This was further assessed in a complete time course up to 15 h (Supplementary Fig. 5A,B). Some late activation was observed but it could not be determined whether this is due to inactivation of the drug during the time lapse microscopy or activation of alternative translational activation pathways. MAPK inhibitors had some effects (Fig. 5D) but these were not as profound as those induced by CDK1 inhibition (Supplementary Fig. 5E-H).

**Figure 4.**
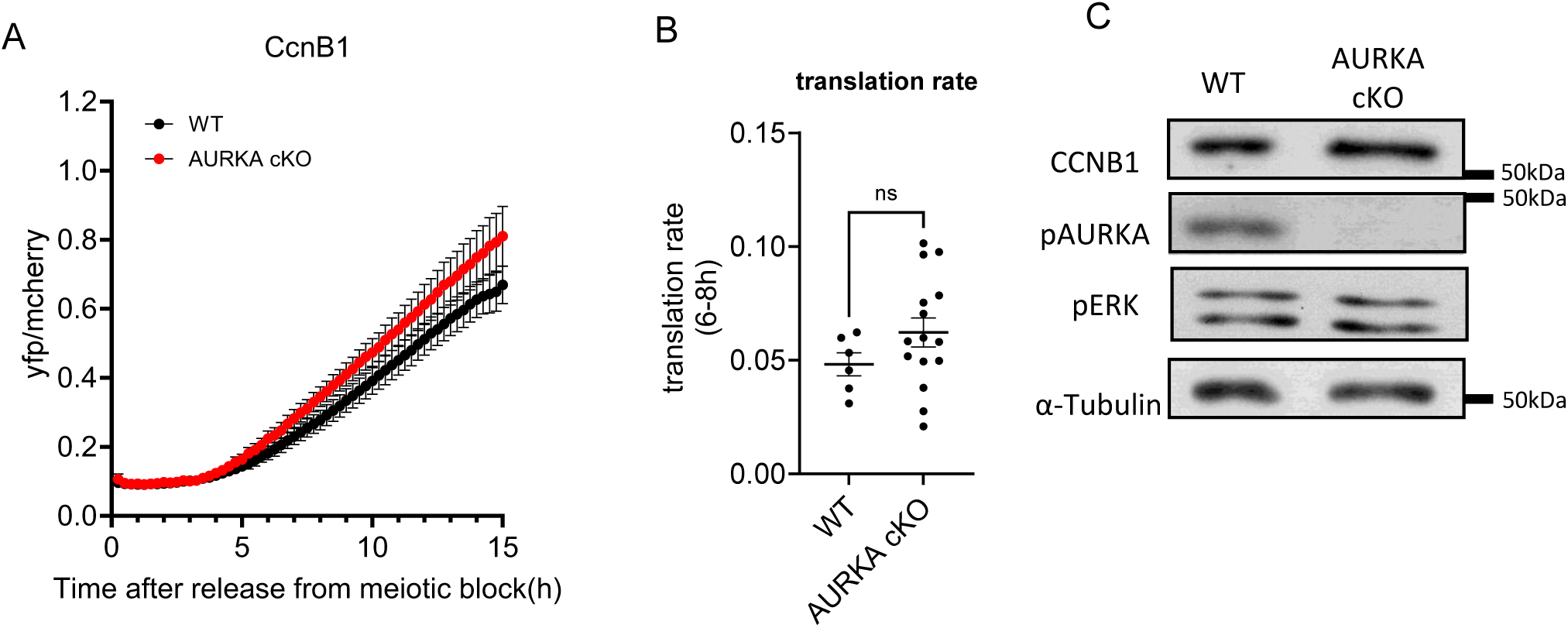
Loss of function of Aurora Kinase A does not compromise *Ccnb1* mRNA translation and endogenous CCNB1 protein accumulation. GV-arrested oocytes were injected with *CyclinB1 (Ccnb1)* reporter and polyadenylated *mCherry* mRNAs. After overnight incubation, oocytes were released in PDE inhibitor (cilostamide)-free medium. YFP and mCherry signals were recorded during maturation by time-lapse microscopy every 15 min for 20 h. (A) The YFP/ mCherry signal ratio for each oocyte was plotted and data are shown as the mean ± SEM. (B) Translation rates were calculated for each oocyte by linear regression of the reporter data between 6 and 8 h after incubation. Two-tailed unpaired Student’s test was used to evaluate statistical significance. ns; not significant. (C) Western blot was performed on lysates of 50 oocytes from wild type (WT) and Aurora Kinase A(AURKA) cKO mouse after 7 h of maturation. Alpha-Tubulin was used as a loading control. The intensity of the endogenous CCNB1 protein was not significantly different.

**Figure 5.**
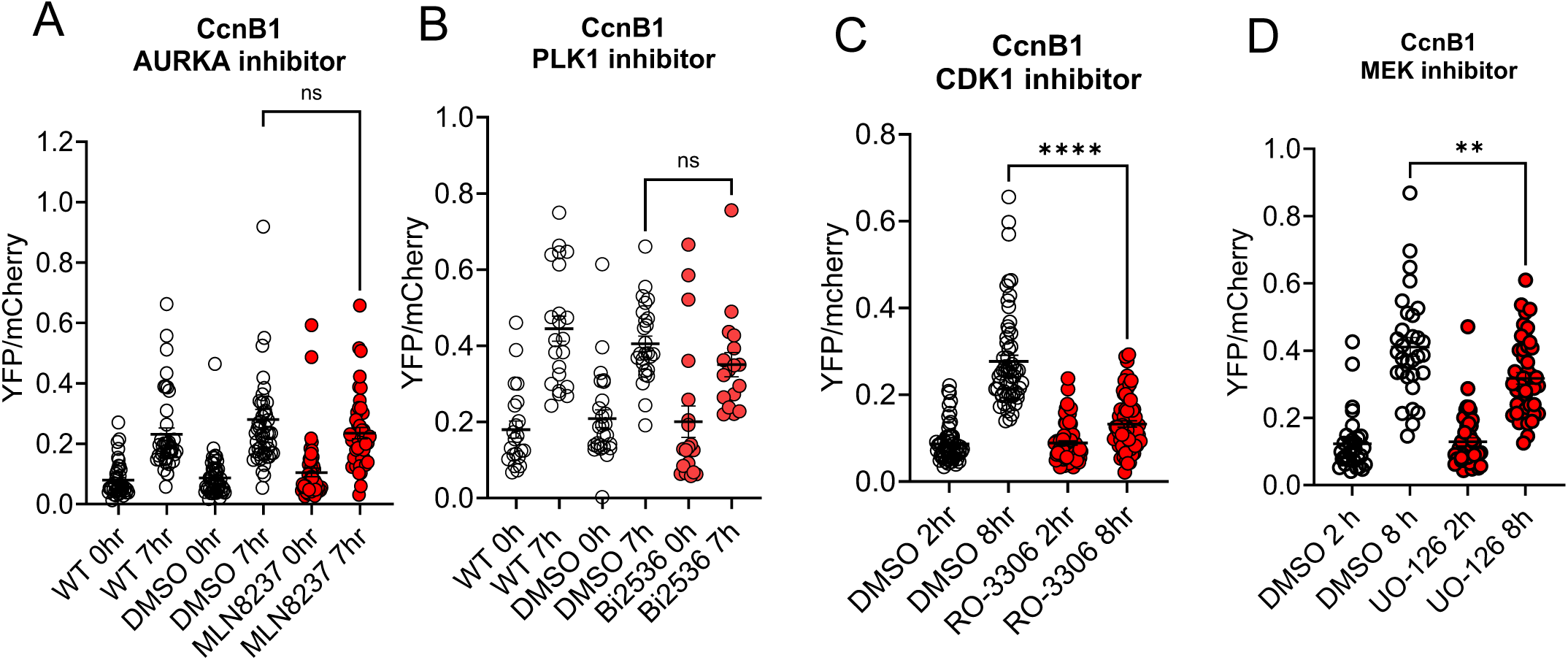
Effect of different inhibitors on *Ccnb1*. GV-arrested oocytes were injected with *CyclinB1 (Ccnb1)* reporter and polyadenylated *mCherry*. (A, B) For MLN8237 and Bi2536 experiments, microinjected oocytes were pre-incubated for 1 h with inhibitors (final concentrations: MLN8237 1µM, Bi2536 0.1µM) before releasing the oocytes into PDE inhibitor (cilostamide)-free medium containing the drugs. (C, D) For the RO-3306 (10µM) and UO-126 (20µM) experiments, oocytes were injected and released into cilostamide-free medium to undergo nuclear envelope breakdown (GVBD). Oocytes that underwent GVBD were collected and transferred to a medium including the above drug. In all experiments, YFP and mCherry signals were plotted at 0 or 2 h incubation and 7-8 h. The data are the mean ± SEM from three different experiments with each dot representing one oocyte. Two-tailed unpaired Student’s test was used to evaluate statistical significance (ns; not significant, **P<0.01, ****P<0.0001). AURKA: Aurora Kinase A, WT: wild type.

To confirm this divergent effect of the two pathways on translation of the reporter, we measured the levels of endogenous CCNB1 accumulation (Fig. 6). CCNB1 protein levels were consistent with the reporter assay, whereby CDK1 inhibition had the most profound effects, followed by MAPK inhibitors: AURKA or PLK1 inhibition had no or minor effects on CCNB1 accumulation in MI (Fig. 4C, 6). Therefore, translation of the endogenous mRNA and of a reporter consistently show that the AURKA/PLK1 pathway has no effect on CCNB1 translation, regardless of CPEB1 destabilizing effect.

**Figure 6.**
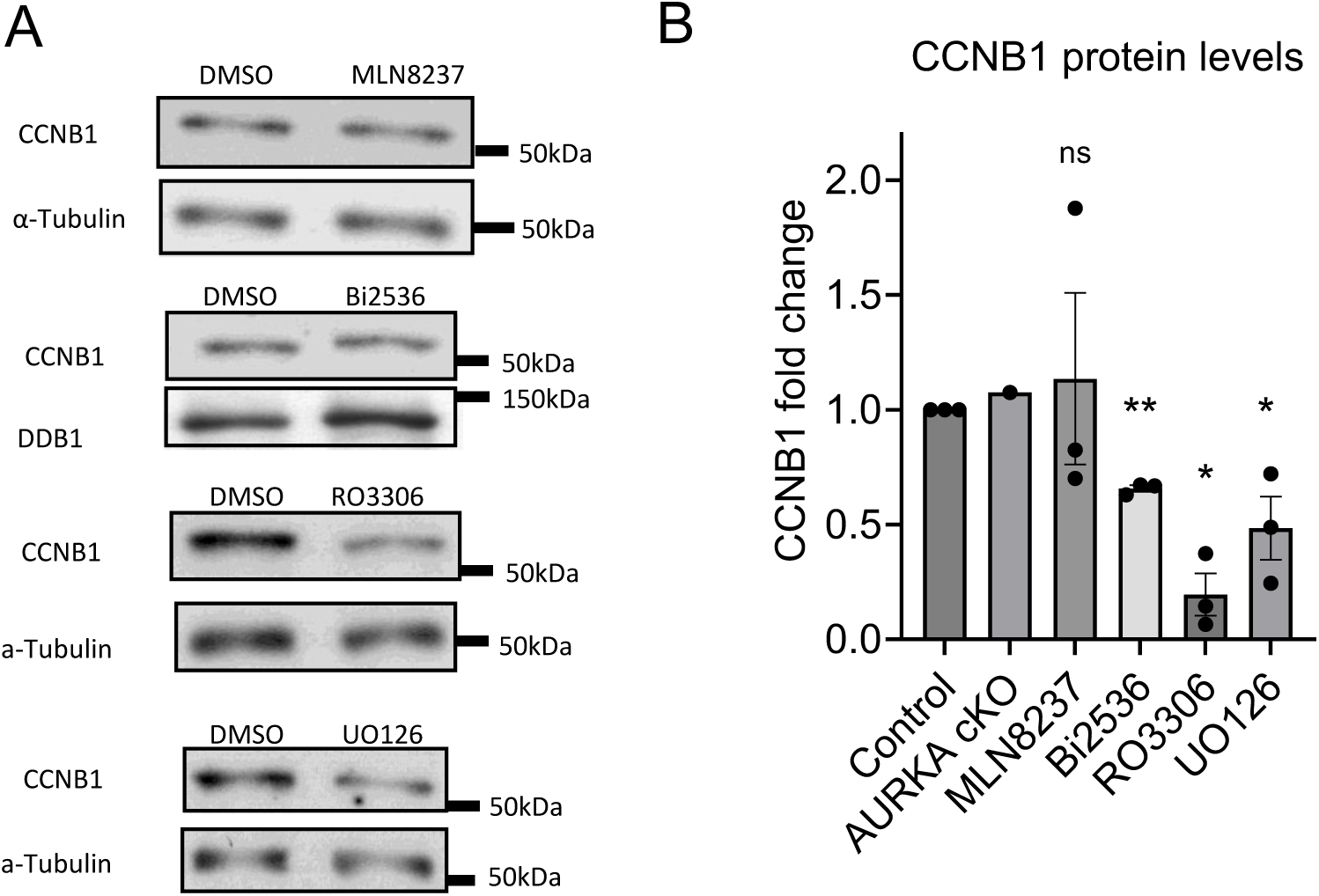
Endogenous CCNB1 protein accumulation with different inhibitors. (A) CyclinB1 protein (CCNB1) accumulation in MI oocytes (7 h incubation) with different treatments. α-Tubulin or DDB1 was used as a loading control. 50-75 oocytes per lane were loaded for each experiment. (B) Quantification of western blot run in the above conditions. The data are CCNB1/ α-Tubulin or DDB1 ratios and are expressed as fold changes over DMSO control. Each bar is the mean ± SEM of three independent biological. Two-tailed unpaired Student’s test was used to evaluate the statistical significance between the control and each treatment (ns; not significant, *P<0.05, **P<0.01). AURKA: Aurora Kinase A, WT: wild type.

### Mos translation is not affected by suppression of the AURKA/PLK1 pathway

To verify whether the divergent effect of the two pathways is exclusive to *Cyclin B1* mRNA translation, we repeated the translation experiments with a *Mos* reporter (Fig. 7). Again, we found that only CDK1/MAPK inhibition had a significant effect on *M*os reporter translation.

**Figure 7.**
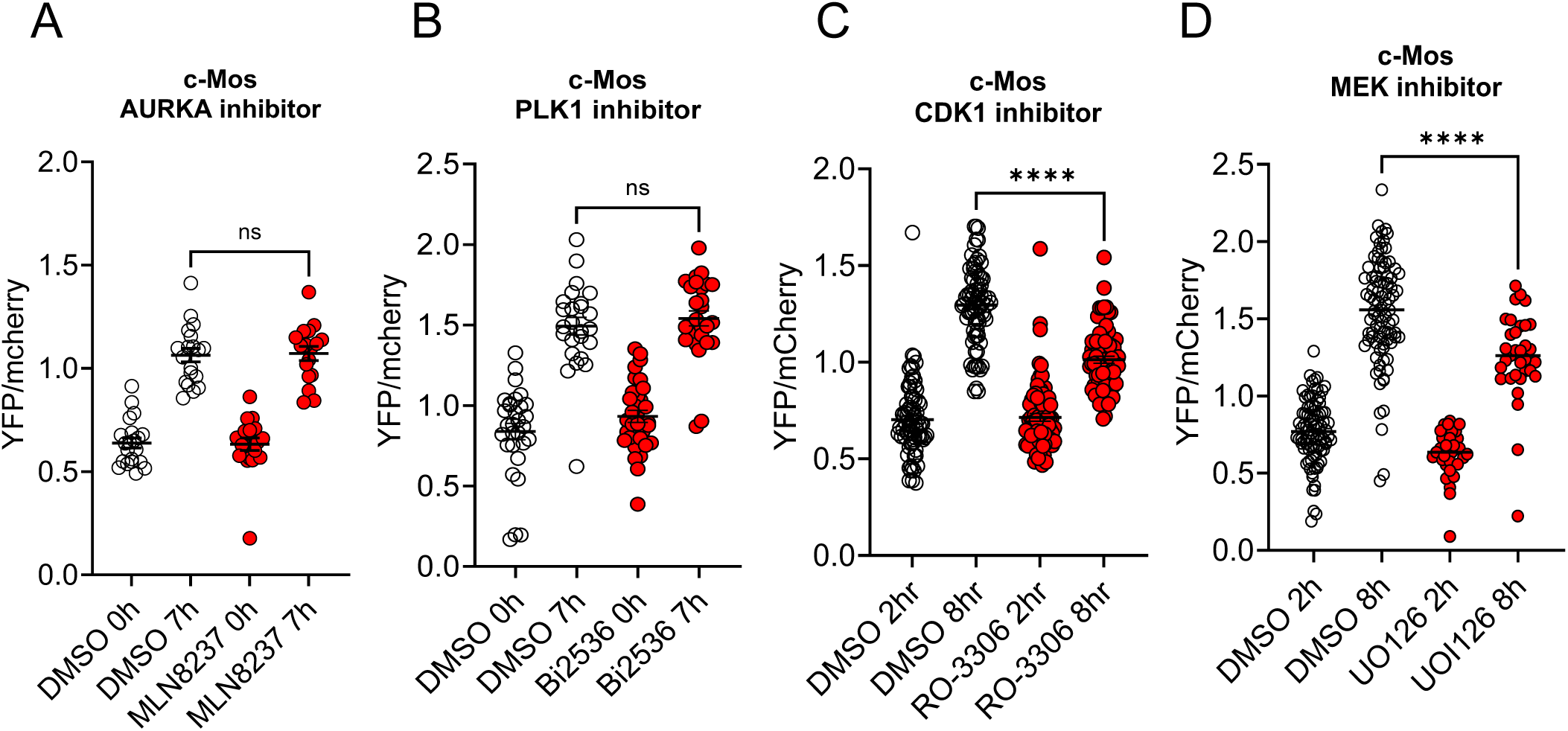
Translation of *Mos* reporter in oocytes treated with different inhibitors. GV-arrested oocytes were injected with *Mos* reporter and polyadenylated *mCherry* mRNAs. (A, B) For the experiments with MLN8237 (1µM) or Bi2536 (0.1µM), microinjected oocytes were pre-incubated for 1 h with inhibitors before releasing the oocytes to PDE inhibitor (cilostamide)-free medium containing the inhibitor. (C, D) For the RO-3306 (10µM) and UO-126 (20µM) experiments, oocytes were injected and released to a cilostamide-free medium to undergo nuclear envelope breakdown (GVBD). Oocytes that underwent GVBD were collected and transferred to the inhibitor-containing medium. In all experiments, YFP and mCherry signals were plotted at 0 h or 2 h incubation (the time when the oocytes were transferred to cilostamide-free drug-containing medium) and 7-8 h of incubation. The data are plotted as the mean ± SEM of the rate of reporter accumulation. Two-tailed unpaired Student’s test was used to evaluate statistical significance (ns; not significant, ****P<0.0001). AURKA: Aurora Kinase A.

Further analysis of the translation of *Mos* after CDK1 inhibition indicates absence of any activation (Supplementary Fig. 5C,D), with accumulation of the reporter throughout maturation occurring at a constant rate.

## Discussion

Here, we present experimental evidence demonstrating the convergence of the AURKA/PLK1 and CDK1/MAPK signaling pathways on phosphorylation of the RNA Binding Protein CPEB1. However, only the CDK1/MAPK pathway is essential for the translational activation of an endogenous mRNA or the translation of a microinjected reporter during the transition from prophase to metaphase I (MI) in mouse oocytes. Consistent with previous findings, we show that the hyper-phosphorylation of CPEB1 leads to its destabilization and degradation. However, under conditions identified in this study, preventing the destabilization induced by AURKA/PLK1 phosphorylation does not perturb the translational activation of these mRNAs during MI. Collectively, all our observations support the overarching hypothesis that the initiation of translation in mouse oocytes primarily relies on CDK1-dependent phosphorylation of CPEB1.

This phosphorylation likely triggers polyadenylation and enhances the translation rate through ribosome recruitment up to MI. This CDK1-dependent phosphorylation is distinct from those induced by the AURKA/PLK1-dependent pathway. In a subsequent phase, both pathways induce hyper-phosphorylation and destabilization of CPEB1. While our findings challenge the notion that CPEB1 destabilization is an indispensable event for the initial translational activation, it remains possible that destabilization and altered stoichiometry of CPEB1 play a role in the translational regulation of a specific subset of maternal mRNAs during the MI-MII transition.

Detailed quantification of the mobility of CPEB1 protein in SDS-PAGE supports the view that early phosphorylation of this protein occurs at the GV/GVBD transition between 1 and 2 h from the release of the meiotic block, or perhaps even earlier with a timing that overlaps with the increase in CDK1 activity. This early step is followed by a further mobility shift between 2 and 4 h. This hyper-phosphorylation is then followed by an overall decrease in CPEB1 protein starting at 6 h (Fig. 8). Even if treatment is delayed after GVBD, CDK1 inhibition with the specific compound RO3306 completely prevents any of the CPEB1 mobility shifts indicating that activation of this kinase is indispensable for both phosphorylation and hyper-phosphorylation. CDK1 inhibition is also the condition that produces the most profound effects on translation of both the endogenous mRNAs and of injected reporters. These findings further confirm that this kinase is at the apex of the molecular cascade orchestrating translational regulations in the oocyte at meiotic reentry.

**Fig. 8.**
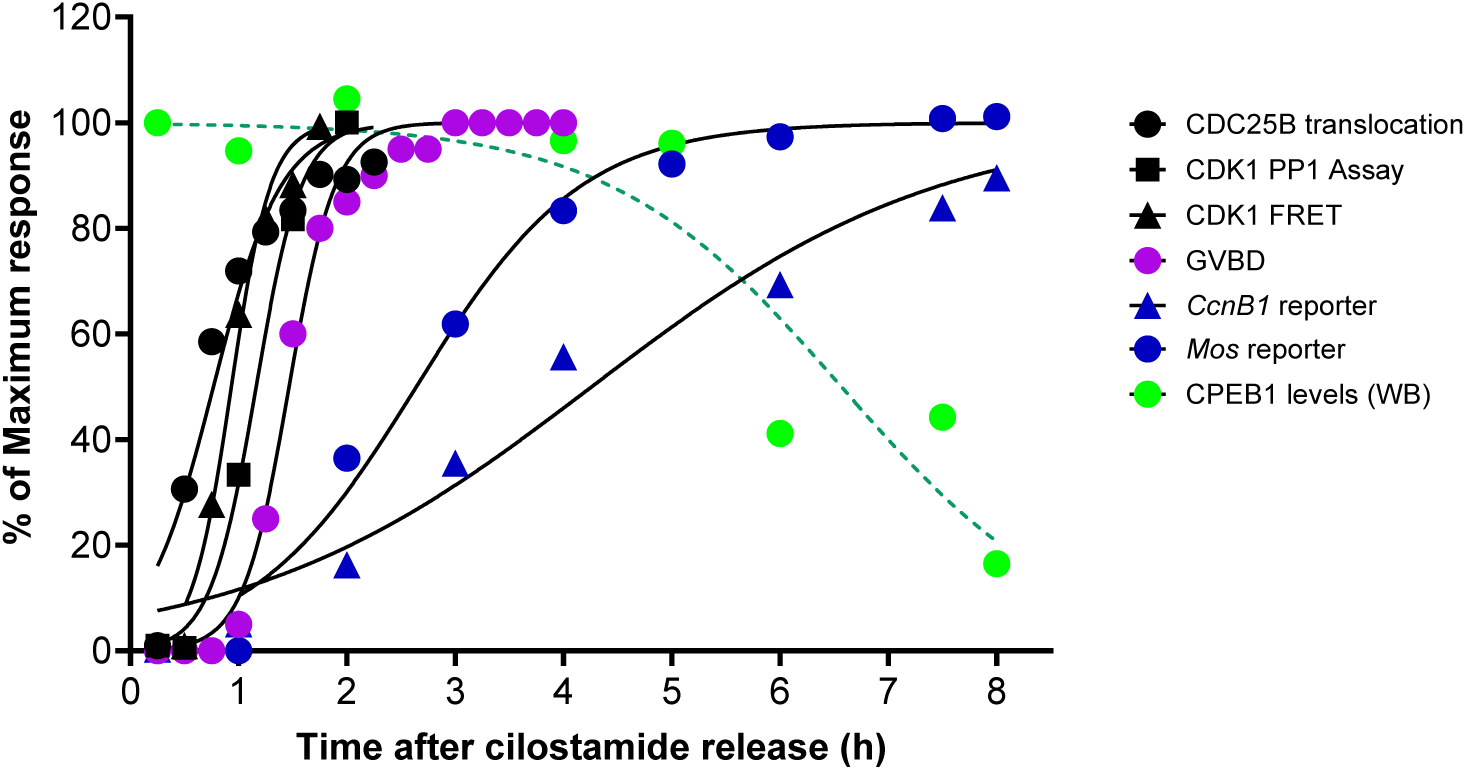
Summary of the timing of landmark events of mouse oocyte in vitro maturation to Metaphase I. The CDC25B protein translocation into the nucleus, which is a proxy of inactivation of the CAMP/PKA pathway, was followed in oocytes microinjected of a fluorescent tagged CDC25B. The CDK1 activity was measured in live oocytes using a FRET reporter or by measuring CDK1 activity in oocyte lysates by monitoring the phosphorylation of a PP1 peptide substrate. The timing of CDK1 activation using the two measurements is comparable. Nuclear envelope breakdown (GVBD) was monitored by time-lapse microscopy and is the average of measurements of more than 100 oocytes. The change in rate accumulation of the *Mos* and *CcnB1* reporter during oocyte maturation was followed by time-lapse microscopy after microinjection of mRNAs coding for the reporters fused to the corresponding 3’UTR. CPEB1 protein levels was assessed in extracts from oocytes cultured for different times after removal of the meiotic block. Each point is the mean of four independent experiments. All data are normalized to the maximum value and reported as % of maximum. In all cases, the standard error of the mean is omitted to improve clarity. Lines are the best fit of the experimental data using a four-parameters logistic equation and variable slope. Average time in Hours +/- 95 Confidence limit: CDC25B = 0.7674 (0.6199 to 0.9234); CDK1 activity measured by PP1 in vitro phosphorylation = 1.161 (1.092 to 1.232); CDK1 activity measured by FRET in live oocytes = 0.9326 (0.8460 to 1.023); GVBD= 1.466 (1.417 to 1.517); *Mos* reporter accumulation = 2.640 (2.360 to 2.928); *CcnB1* reporter accumulation = 4.260 (3.660 to 4.898); CPEB1 levels = 6.566 (5.701 to 7.562). Of note the rate of translation of the reporters is consistent with the timing of increased ribosome loading (see Ref 47).

Our genetic data show that the activation of translation of both *CcnB1* and *Mos* mRNAs is dependent on CPEB1. However, detailed analyses of the time courses of translational activation document significant temporal differences, with *Mos* translation preceding that of *CcnB1*. Inspection of these two mRNAs shows that the cis-acting elements configuration in the 3’ UTR is similar but not identical to that described in *Xenopus* (Stebbins-Boaz et al., 1996, Mendez et al., 2002). The *Mos* 3’ UTR includes a putative *Pum1* (TGTAGATAA), a *Zar1* (TTTGTGT) and a long polyU stretch described as an embryonic CPE. Conversely, at least four functional CPEs have been mapped in the 3’ UTR of *CcnB1*. Therefore, the different time course in mouse oocytes may be due to the presence of a single CPE in *Mos* and multiple CPEs in *CcnB1*, as proposed for *Xenopus* oocytes. Nevertheless, the presence of other regulatory elements may contribute to the difference in the timing of translation. In *Xenopus* oocytes, an additional RBP, MUSASHI, interacts with the CPE in the 3′ UTR of the *Mos* mRNA and that this interaction is necessary for early *Mos* mRNA translational activation (Charlesworth et al., 2006, Weill et al., 2017, Arumugam et al., 2010). We could not identify a perfect consensus for MUSASHI binding in the 3’UTR of mouse *Mos*. However, this mouse regulatory region has a consensus site for ZAR1 binding (TSC element) (Charlesworth et al., 2012). It remains to be determined whether a *Zar1* interaction contributes to the temporal pattern of translation observed, as proposed in *Xenopus* oocytes (Heim et al., 2022). In addition to a distinct initial timing of activation, translation between MI and MII follows a different pattern for the two mRNAs, with ribosome loading and the rate of translation of *Mos* reaching a plateau around 5-6 h or MI, whereas ribosome loading and translation of the CcnB1 mRNA continues to increase between MI and MII. This difference is reminiscent of the early and late translations described in *Xenopus* oocytes.

A detailed analysis of the timing of the different events triggered during oocyte maturation strongly suggest that CDK1 activation precedes translational activation for both the *Mos* and *CcnB1* mRNAs (Fig. 8). This initial translational activation precedes any detectable destabilization of CPEB1 by 3-4 h. These temporal correlations strongly indicate that the initial ribosome loading, reporter accumulation, and endogenous protein accumulation are due to a critical CDK1-dependent phosphorylation of CPEB1. This phosphorylation triggers translational activation but not destabilization of the protein. Available phosphoproteomics data did not detect phosphorylation of the residue that in the mouse corresponds to Ser174 of *Xenopus* CPEB1.

Therefore, the activating phosphorylation likely corresponds to one of the CDK1 or MAPK sites that have been mapped. Later, during oocyte maturation, destabilization of CPEB1 may be necessary for further translational activation of *CcnB1* at the MI-MII transition. Unlike that reported in *Xenopus*, we could not determine whether translation of *Mos* and *Ccnb1* mRNAs are affected differently by CPEB1 destabilization, even though the temporal patterns described above are consistent with the view that *CcnB1* mRNA further translational activation at the MI-MII transition coincides with significantly decreased CPEB1 levels, whereas *Mos* translation remains unchanged. We could not directly investigate the effect of CPEB1 stabilization on the MI-MII transition because the *Cpeb1^fl/fl^*Zp3-cre and AURKA cKO as well as the pharmacological inhibitors all prevent anaphase I and polar body extrusion. In *Xenopus* oocytes, overexpression of a mutant *Cpeb1* resistant to degradation prevents the MI-MII transition and CCNB1 synthesis at this stage (Mendez et al., 2002). Conversely, overexpression of a stable CPEB1 only partially prevents the MI-MII transition in mouse oocytes (Sha et al., 2017).

Consistent with the hypothesis proposed in previous reports, we conclude that hyper-phosphorylation of CPEB1 is the signal triggering destabilization. The time course of the appearance of the hyper-phosphorylated CPEB1 form overlap with the timing of decreased levels of CPEB1. In addition, any condition that delays or prevents hyper-phosphorylation also prevents destabilization. The only observation inconsistent with this hypothesis is that MAPK inhibition causes CPEB1 stabilization while in the hyper-phosphorylated state. One explanation that reconciles this discrepancy is the possibility that MAPK inhibition interferes with mechanisms required for proteasome activation and ubiquitination. Further experiments are required to test this possibility. Moreover, post-translational destabilization of CPEB1 may not be the only mechanism leading to a decrease CPEB1 protein. It should be noted that we have shown that ribosome loading onto *Cpeb1* mRNA decreases with a time course not dissimilar to that shown for the CPEB1 decline (Luong et al., 2020). Therefore, it is most likely that the decline in CPEB1 protein observed is caused by a combination of decreased translation and destabilization of the synthesized protein.

The role of the AURKA-mediated phosphorylation of CPEB1 in translational activation has been controversial (Radford et al., 2008). Our data using the mouse model indicate that AURKA signaling does have a role in CPEB1-regulated translation, being involved in destabilization of this RBP. However, all our data are consistent with the view that activation of this pathway does not contribute to the translational activation, at least up to 8 h of incubation or around metaphase I. Nevertheless, it remains possible that the AURKA-dependent destabilization of CPEB1 plays a role at the MI-MII transition. Because oocytes lacking AURKA do not progress beyond metaphase I, this genetic model does not provide information as to whether AURKA-mediated CPEB1 destabilization plays a role at the MI-MII transition. In a previous report, oocytes lacking the *AURKA* homolog, *Aurkb*, had elevated levels of translation and CPEB1 was destabilized earlier in maturation. When a *Ccnb1* 3’ UTR luciferase reporter was expressed in these *Aurkb* cKO oocytes, translation was also elevated compared to WT oocytes expressing this reporter. Taken together with findings presented here, it is possible that CDK1 activity is increased in *Aurkb* cKO oocytes, a mechanism to examine in future studies. Also in that report, we found that AURKA activity was increased in *Aurkb* cKO oocytes. Consistent with data reported here, CPEB1 phosphorylation was altered, and the protein was stabilized in AURKA cKO oocytes. However, when an *Aurkc* 3’ UTR luciferase reporter was tested, translation was not observed. This result led to the conclusion that AURKA controls CPEB1-dependent translation. This conclusion conflicts with the findings described here. It is possible that the AURKA pathway only controls a sub-set of mRNAs for translation, perhaps dictated by sub-cellular localization or other UTR regulatory sequence elements. AURKA could also have different functions or interactions with CPEB1 during later meiotic maturation time points.

AURKA cKO is associated with major dysfunction of the spindle and microtubule dynamics (Blengini et al., 2021). Therefore, the possibility needs to be considered that localized translations in a compartment around the spindle may be disrupted in this genetic model (Waldron and Yajima, 2020, Pascual et al., 2020).

At present, we could not determine whether the CDK1 effects on translation are direct or entirely mediated by activation of MAPK. MAPK activation is in large part dependent on CDK1 activity, and the synthesis of the upstream kinase MOS. MAPK activation measured by phosphorylation becomes detectable after 2-4 h of meiotic resumption*. Mos* mRNA translational activation is detected around the same time and corresponding protein accumulation is detected between 2 and 4 h. Given the resolving power of our measurements, we cannot determine whether the timing of this activation is compatible with the possibility that all the CDK1 effects on translation are indirect and mediated entirely by MOS/MAPK activation. It should be noted that the rate of translation of *Mos* mRNA is high in comparison for instance to that of *CcnB1*. This difference may indicate considerable rate of synthesis of MOS protein also in GV oocytes. Since, the protein is non-detectable by western blot in these times, one could argue that newly synthesized protein is rapidly degraded likely because there is no sufficient CDK1 activity to stabilize it. If this hypothesis is correct, then it is possible that at the beginning of cell cycle reentry, MOS becomes stabilized by CDK1 phosphorylation and that this stabilization allows some MAPK activation prior to activation of *Mos* mRNA translation. Further experiments are necessary to address this issue.

In summary, our data support the hypothesis that both the CDK1/MAPK and AURKA/PLK1 pathways are involved in CPEB1 regulation as they converge on the phosphorylation of this protein. In regulating the phosphorylation of this RBP, the two pathways have a role in translational regulation during meiotic maturation. Initial CPEB1-mediated translational activations are distal to CDK1 activation but are independent on AURKA/PLK1 mediated phosphorylations. Later during MI, both kinase cascades contribute to the destabilization of CPEB1. This late destabilization in turn may contribute to further translational activation of *CcnB1* mRNA during the transition from MI to MII. It therefore possible that a subset of translational activations at these later stages of meiotic maturation requires the activity of both kinases.

## ACKNOWLEDGMENTS

We thank Dr. Raúl Méndez and Dr. Gonzalo Fernandez-Miranda (The Barcelona Institute of Science and Technology) for sharing the CPEB1-targeted mice. We also thank Dr. Xiaotian Wang and Dr. Nozomi Takahashi for the helpful discussion and technical advice. This work was supported by NIH R01 GM116926, and in part by NIH/NICHD P50 HD055764 Center for Research, Innovation and Training in Reproduction and Infertility to MC, and NIH/NIGMS R35 GM136340 to KS. EMD was supported by a grant “Emergence 2023-2024” from Sorbonne Université. CK is supported by a fellowship from the Japan Society for the Promotion of Science (202112871) and Tokyo University.

## Materials and Methods

### Mice

All procedures involving mice were approved by the University of California, San Francisco’s Institutional Animal Care and Use Committee (Approval # AN197697-00B). Animal care and use followed relevant guidelines and regulations. Mice were housed in a 12-12 h light-dark cycle with access to food and water ad libitum, under constant temperature. All animals used were of the *C57BL/6J* inbred strain. CPEB1-targeted mice were a gift from Dr. Raul Mendez (Calderone et al., 2016) and bred in our laboratory. Aurora kinase A (AURKA) conditional knock-out mice were descrived previously (Blengini et al., 2021) and bred in the Schindler laboratory. Oocytes were shipped from the Schindler lab for western blot processing and mice were shipped for live reporter assay imaging. These mice were approved by the Rutgers University Institutional Animal Care and Use Committee (protocol 201702497).

### Oocyte collection and culture

Female mice (3-4 weeks old) were intraperitoneally injected with 5 IU pregnant mare serum gonadotropin (PMSG; MyBioSource, MBS173236) to induce follicle growth. After 44 h, the ovaries were dissected and cumulus-oocyte-complexes (COCs) were collected in media containing HEPES-modified minimum essential medium Eagle (Sigma-Aldrich, M2645) supplemented with 6 mM sodium bicarbonate (JT-Baker, 3506-1), 0.2 mM sodium pyruvate (Gibco, 11360070), 75 µg/ml Penicillin+10 Vg/ml Streptomycin (Genesee Scientific Corporation (GenClone), 25-512), 3 mg/ml BSA (Sigma-Aldrich, SIAL-A3311), and a PDE inhibitor (1 µM cilostamide (Calbiochem, 231085)). Using a glass pipette, aspiration was performed to remove the cumulus cells surrounding the eggs. Denuded oocytes were maintained at the GV stage in alpha-minimal essential medium (α-MEM) (Gibco) supplemented with 0.2 mM sodium pyruvate, penicillin–streptomycin, and 1 µM cilostamide at 37°C under 5% CO2. After microinjection and overnight incubation, oocytes were transferred to α-MEM without cilostamide and allowed to mature. For GV state oocyte sample, oocytes were kept in cilostamide medium. For AURKA oocytes sample collection, oocytes were kept in MEM containing 2.5µM milrinone (Sigma-Aldrich, M4659). Then oocytes were transferred to MEM without milrinone and allowed to mature. Oocytes were then collected at various time points. For AURKA/PLK1 pathway experiments, oocytes were preincubated for 1 hour with 1 µM MLN8237 (Selleckchem, s1133) or 0.1 µM Bi2536 (VWR, 101757-124) before release from cilostamide. For 10 µM RO3306 (Selleckchem, s7747) and 20 µM UO126 (Selleckchem, s1102) experiments, the inhibitors were added after 2 h incubation without cilostamide. The inhibitor experiments were done without oil cover.

### Construction of fluorescent protein reporters

The 3’UTR sequences of *Ccnb1* and *Mos* were obtained by amplification of mRNA from oocytes with appropriate primers (Supplementary Method) and fused with Ypet ORF. (C483S)-CDC25B coding sequences were also fused with Ypet ORF. The CDK1 FRET sensor was a gift from Jonathon Pines (Addgene plasmid # 26064). They were subcloned in the pCDNA 3.1 vector containing a T7 promoter and the constancy was confirmed via DNA sequencing. Linearized cDNAs were then transcribed *in vitro* to synthesize mRNAs with mMESSAGE mMACHINE T7 Transcription Kit (Ambion, AM1344) and purified using MEGAclear kit (Ambion, AM1908). The *mCherry* reporter was similarly produced and polyadenylated (150-200nts) using Poly(A) Tailing kit (Ambion, AM1350).

### Microinjection and Time-lapse microscopy

Collected GV-arrested oocytes were injected with 5-10 pl of 12.5 ng/µl solution of the Ypet reporters along with a *mCherry* reporter or FRET reporter using a FemtoJet Express programmable microinjector with an Automated Leica microinjection Microscope System (LEICA DMI 4000 B). After overnight pre-incubation in α-MEM with 1 µM cilostamide, oocytes were released from cilostamide and matured *in vitro* under time-lapse microscopy. Live cell imaging was performed under a Nikon Eclipse T2000-E equipped with a mobile stage and an environmental chamber at 37°C and 5% CO2. Filter set: dichroic mirror YFP/ CFP/ mCherry 69008BS; Ypet channel (Ex: s500/20x 49057 Em: D535/30m 4728811); mCherry channel (Ex: s580/25x 49829 Em: D632/60m); Cerulean channel (Ex: 430/25x 49829 Em: 480/40m 49287), for FRET channel (Ex: 430/25x 49829 Em: D535/30m 47281). Images were processed and fluorescence was quantified using MetaMorph (Molecular Devices). For reporter assay, YFP and mCherry channels were corrected by background. Then, the ratio of Ypet fluorescence and the maximum level of mCherry signal was plotted as an accumulation of translation activity. For the FRET assay, YFP, CFP and FRET channels were corrected by background. FRET channel was corrected for the YFP bleed-trough, and FRET was calculated as FRET channel/CFP channel.

### Western Blot Analysis

Oocytes at indicated times of incubation under maturing conditions were collected in 0.1% w/v PBS/PVP and mixed with Laemmli sample buffer supplemented with β-mercaptoethanol, proteinase inhibitor and phosphatase inhibitor. After boiling at 95°C for 5 min, the lysates were separated on 8% v/v or 12% v/v polyacrylamide gels and transferred to a Polyvinylidene difluoride (PVDF) membrane (Millipore ISEQ00010). Membranes were incubated in 5% w/v milk buffer for 1 h at room temperature and incubated in primary antibody overnight at 4°C. The antibodies used were as follows: CPEB1 (Abcam, ab73287, 1:1000), Cyclin B1 (Cell-Signaling, 4138, 1:1000), Aurora kinase (Cell-Signaling, 2914T, 1:500), MAPK (Cell-Signaling, 4377, 1:1000), α-tubulin (Sigma-Aldrich, T6074, 1:10,000), DDB1(Abcam, ab109027, 1:4000), and T320-PP1 (Abcam, ab62334, 1:30,000). Membranes were washed in 1 x TBST and incubated in the appropriate secondary antibodies (anti-Rabbit, GE Healthcare, NA934V; anti-mouse, GE Healthcare, NA931V) for 2 h at room temperature. Clarity Western ECL substrate (Bio-Rad, 170-5061) was used to develop the membranes, and ImageJ 1.53a was used for quantification for the signals.

### In Vitro Cdk1 assay (PP1 Assay)

30 oocytes were collected in 30 µl of 2x Kinase buffer (100 mM Hepes, 30 mM MgCl2, 2 mM EGTA, 10 mM CaCl2, 2 mM DTT, 2 µg/ml Leupeptin, 2 µg/ml Aprotinin, and 2 µM okadaic acid). The oocytes were then lysed by freeze-thawing them three times in liquid nitrogen.

Subsequently, 0.1 mM ATP, 10 mM DTT, and 2 µg of recombinant peptide PP1-GST were added to the lysates as the substrate. The lysates were incubated at 30°C for 30 minutes, and the reactions were stopped by boiling them at 95°C for 5 minutes after adding Laemmli sample buffer (Bio-Rad, 161-0747) supplemented with β-mercaptoethanol (Gibco 21985023). Finally, CDK1 activity was assessed by measuring the phosphorylated T320 of the total substrate using western blot analysis.

### Statistical analysis

All data are shown as mean ± SEM. All statistical analyses were performed using the Graph Pad Prism 9 software. Student’s t-tests and Mann-Whitney tests were performed and a P value < 0.05 was considered statistically significant.

