## Supplementary material and figures for "Multiple intersecting pathways are involved in the phosphorylation of CPEB1 to activate translation during mouse oocyte meiosis"

| Name | Prime (5'-3') |
|---|---|
| *Ccnb1* FW | CACCATCACCATTGACTCCAATAGAC |
| *Ccnb1* Rev | GATCAGCGGGTTTAAACAAGCTTTCC |
| *Mos* FW | CAATAATTCTAGACTCCATCGAGCCGATGTAGAG |
| *Mos* Rev | CCACCTGGATCCGAAGTTCGTGGTAACTTTATTTC |
| *mCherry* FW | GAACGGCCACGAGTTCGAGA |
| *mCherry* Rev | CTTGGAGCCGTACATGAACTGAGG |
| FRET FW | TTGGAGCGCTTGACCTTGGGCTAAGGATCCACCGGATCTAGATAACTGATCATAAT |
| FRET Rev | TCGGCATGGACGAGCTGTACAAGGGCGGCGGCTTGCCACCATTGGAGCGC |
| Ypet Rev | GATCCGGTGGATCCTAAAGATCTCTTATAGAGCTCGTTC |
| T7 FW | GAGAACCCACTGCTTAC |
| Vector FW | CTTGACGAGTTCTTCTGAGCGGGAC |
| Vector Rev | GTCCCGCTCAGAAGAACTCGTCAAG |

**Primer sequences used for reporter assay.**

### Supplementary Figures

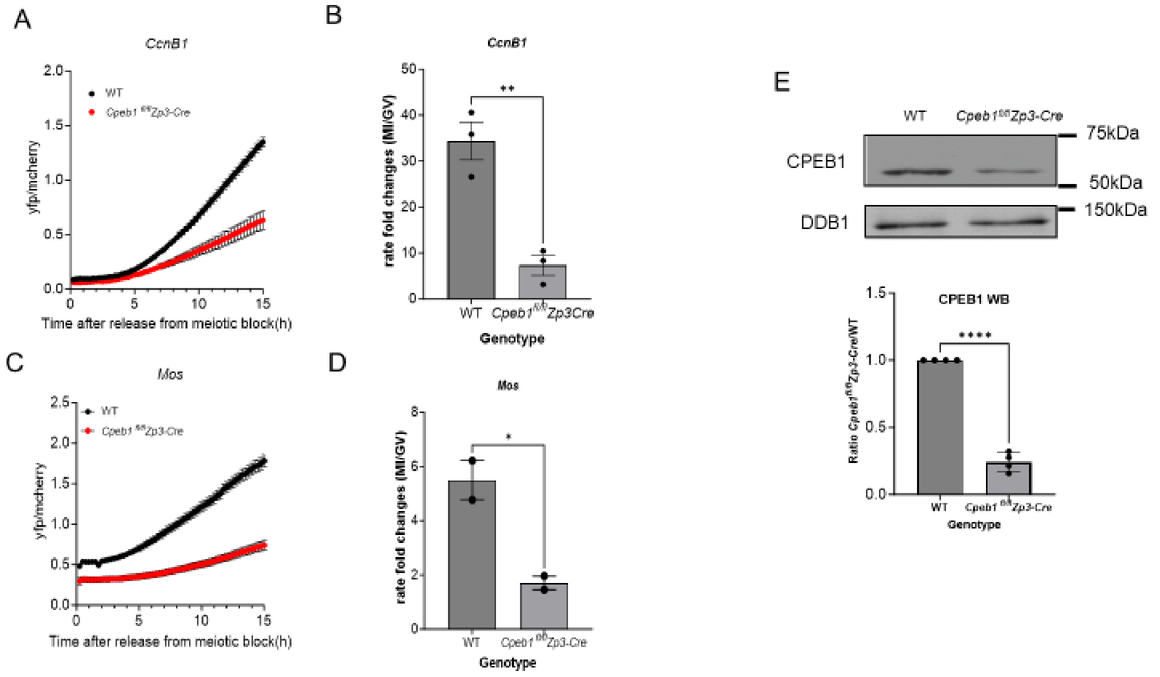


**Supplementary Fig. 1 CPEB1 plays a major role in translational activation**

(A-D) GV-arrested oocytes were injected with *CyclinB1 (Ccnb1)* or *Mos* reporter and polyadenylated *mCherry*. After overnight incubation with 1μM cilostamide (PDE inhibitor), oocytes were released from cilostamide and matured. YFP and mCherry signals were recorded by time-lapse microscopy every 15 min for 20 h. The YFP/ mCherry signal ratio for each oocyte was plotted and data are shown as the mean ± SEM. The value is shown as a fold change. The bar graph shows the ratio of translation rate between GV and MI stages. Each bar is the mean ± SEM of three or more independent biological replicates. Two-tailed unpaired Student's test was used to evaluate statistical significance (*P<0.05, **P<0.01). (E) Residual CPEB1 expression in CPEB1$^{fl/fl}$Zp3-cre mice. Western blot was performed on lysates of 30 oocytes from wild-type (WT) and CPEB1$^{fl/fl}$Zp3-cre mice. A graph reports the quantification of the western blot from four independent biological replicates. The data are the mean ± SEM of the CPEB1/DDB1 ratios plotted as the changes over the control. Two-tailed unpaired Student's test was used to evaluate statistical significance (****P<0.0001).

A

**Potential phosphorylation sites in mouse CPEB1**

| *Position in the sequence* | *Peptide sequence* | *Fold change MI/GV* | *Notes* |
|---|---|---|---|
| S4 | MAF**S**LEEAAGRIKDCWDNQ | Not detected in GV | Unknown |
| S43 | SNANIFRRINAILDD**S**LDFSKVCTTPINRGI | Not detected in GV | Potential PLK1 substrate |
| S207 | SGSDHLSDLISSLRI**S**PPLPFLSMTGNGPRD | Not detected in GV | Potential MAPK* |
| S245 | SRMDQEQAALAAVAP**S**PTSAPKRWPGASVWP | 21.63363 | Potential MAPK |
| S85 | ETVTSRMLFPTSAQE**S**PRGLPDANGLCLGLQ | 6.883888 | Potential Cdk1 |
| S181 | GSRLDTRPILDSRSS**S**PSDSDTSGFSSGSDH | 0.498547 | Potential MAPK* |
| T52 | NAILDDSLDFSKVCT**T**PINRGIHDQLPDFQD | Not detected in GV | Found among PLK1 and AurkA substrates |

B

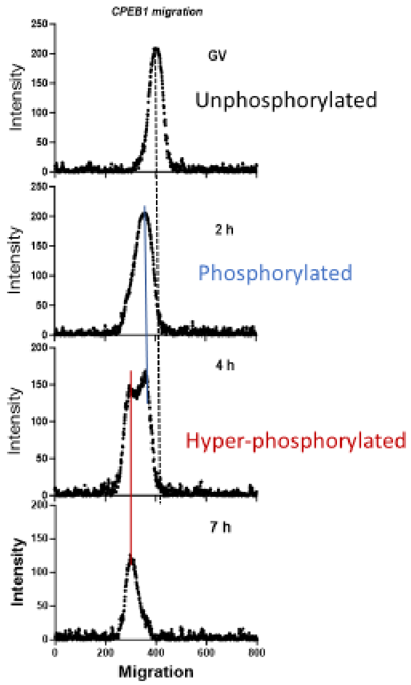


**Supplementary Fig. 2 Multiple CPEB1 phosphorylation and the site in mouse**

(A) The table shows the potential phosphorylation sites in mouse CPEB1 by phosphoproteome approach (ref 27). Residue with the highest probability of being phosphorylated is reported in red. (B)The image shows the time shift of CPEB1 migration using the intensity of the western blot band. Western blot was performed on lysates of 30 oocytes from wild-type (WT). The left side of the x-axis shows the intensity of the upper band (hyper-/ phosphorylation).

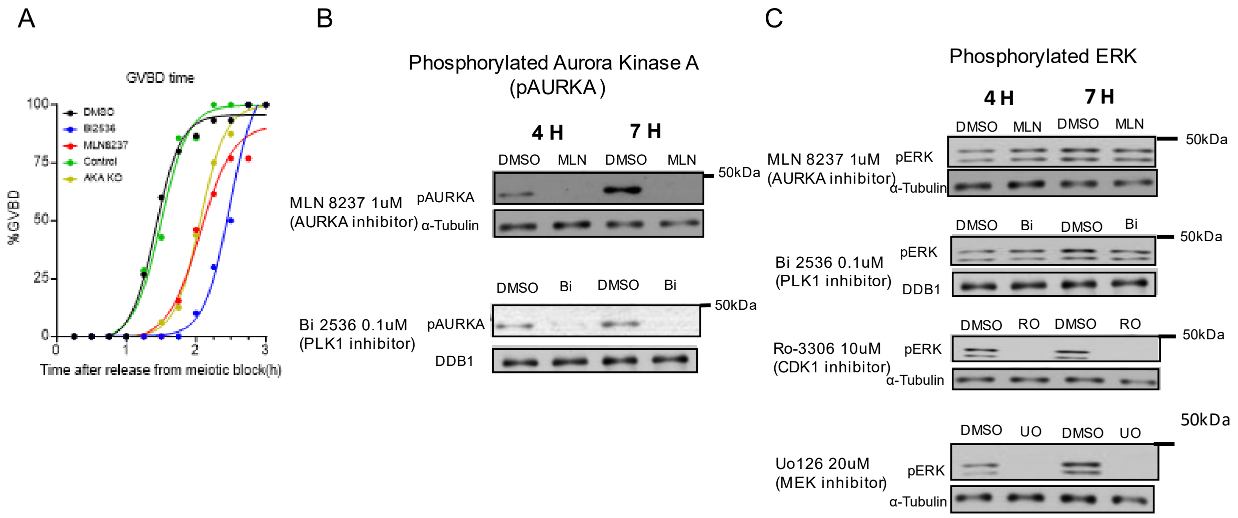


**Supplementary Fig. 3 Delay in GVBD time and the efficiency of different treatments**

(A) Time-lapse analysis of the timing of GVBD in different treatments. (B) Representative western blot image showing the efficiency of MLN8237 and Bi2536 treatment in phosphorylation of Aurora Kinase A (AURKA). (C) Representative western blot images showing the efficiency of different treatments in phosphorylation of ERK1/2. Alpha-Tubulin or DDB1 was used as a loading control. Western blot analysis was conducted on lysates of 30 oocytes.

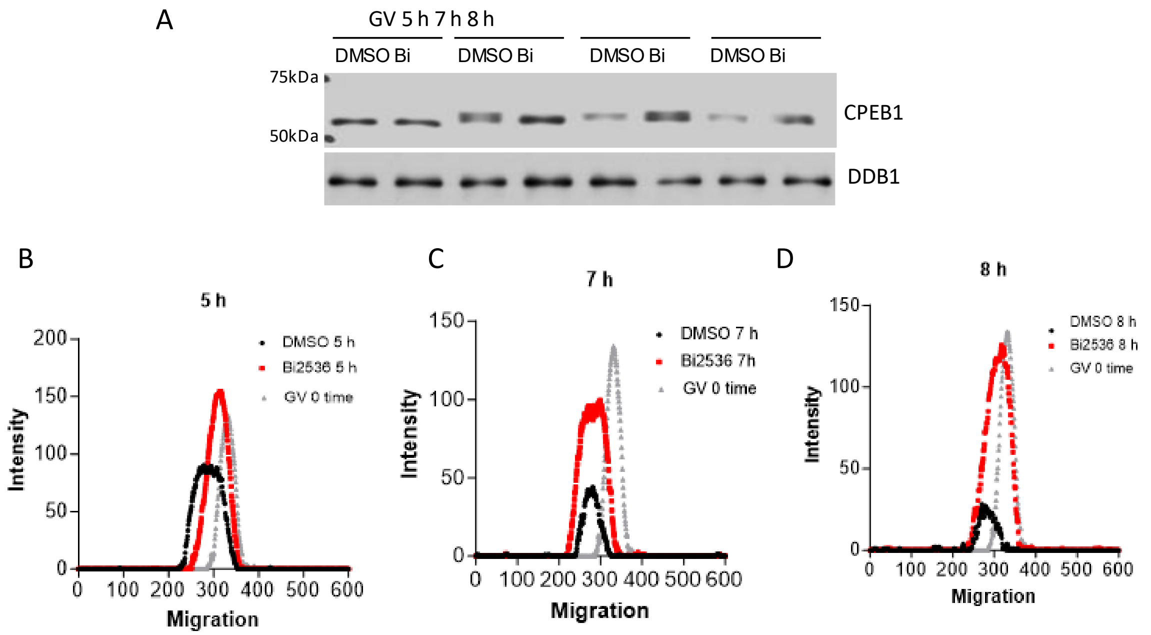


**Supplementary Fig. 4 Inhibition of PLK1 stabilizes phosphorylated CPEB1**

(A) A representative western blot of time course of CPEB1 phosphorylation with Bi2536 treatment (30 oocytes/ lane). DDB1 was used as a loading control. (B) Intensity graph of western blot band showing the migration of CPEB1 phosphorylation during oocyte maturation in oocytes treated with Bi2536. Bi2536 treatment group delays the phosphorylation and CPEB1 is stabilized in 7 and 8 h.

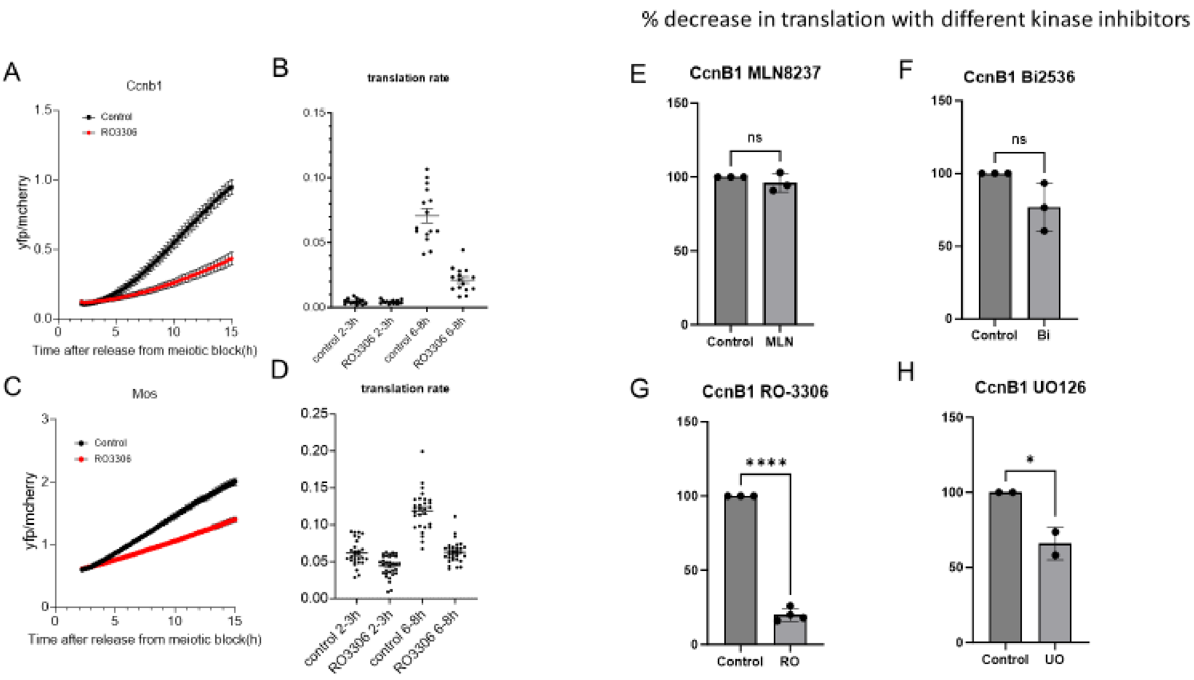


**Supplementary Fig. 5 CDK1 inhibition causes decrease in translation of *Ccnb1* and *Mos***

(A-D) GV-arrested oocytes were injected with *CyclinB1 (Ccnb1)* or *Mos* reporter and polyadenylated *mCherry*. After overnight incubation with 1µM cilostamide (PDE inhibitor), oocytes were released from cilostamide and matured. YFP and mCherry signals were recorded by time-lapse microscopy every 15 min for 20 h. The YFP/ mCherry signal ratio for each oocyte was plotted. The translation rate for each oocyte was calculated by linear regression of the reporter data with an indicated window. All data are shown as the mean ± SEM. (E-H) The graphs show the % decrease in translation of *Ccnb1* with different kinase inhibitors referred to control. The data are plotted with the mean ± SEM and each point represents a different experiment and biological replicate. Two-tailed unpaired Student's test was used to evaluate the statistical significance (ns; not significant, *P<0.05, ****P<0.0001).
